# A Two-Phase Fluid Structure Interaction Model of Mucociliary Clearance Driven by Cilium

**DOI:** 10.1101/2025.05.07.652624

**Authors:** Kavin Vishnu, Karupppasamy Subburaj, Monika Colombo

## Abstract

This study introduces a two-phase fluid-structure interaction (FSI) model of mucociliary clearance (MCC), utilizing direct cilia modeling and non-Newtonian Carreu rheology for the mucus layer (ML). A novel method for prescribing cilia beat patterns (CBP) was developed, enabling realistic motion replication by imposing displacement at specific cilium points. Computational simulations evaluated MCC performance by comparing rheological properties and assessing the interaction between the ML and periciliary layer (PCL). Results highlight improved ML-PCL coupling and enhanced mucus transport using non-Newtonian ML. This approach enables deeper insights into mucus dynamics and the role of cilia motion in MCC.

## 1. Main text introduction

Over the past few decades, computational fluid dynamics (CFD) models of mucociliary clearance (MCC) have significantly advanced, aiming to elucidate phenomena critical to lung health. Many respiratory diseases, such as chronic obstructive pulmonary disease, cystic fibrosis, and asthma, are thought to be directly related to MCC dysfunction (1,2). Recently, CFD models have also been utilized to investigate the behavior of pathogens like SARS-CoV-2, antibodies, and aerosol deposition within the respiratory system (3,4). These models may help improve understanding therapeutic techniques for lung disease, although research into airway clearance techniques for enhancing MCC via CFD modeling remains limited (5). Moreover, CFD models allow for exploring how various parameters, including temperature, humidity, and ciliary abnormalities, influence MCC (6,7).

The driving force of MCC is the coordinated motion of motile cilia, as opposed to primary cilia, which function as biological sensors. The lungs are lined with approximately 3 × 10^12^ motile cilia, each composed of motor protein complexes known as dyneins. These dyneins convert adenosine triphosphate (ATP) into mechanical energy, driving cilia movement (8,9). Motile cilia are immersed in the airway surface liquid (ASL), which consists of two layers: the lower periciliary layer (PCL), which is water-like, and the upper mucus layer (ML), which is highly viscous and exhibits shear-thinning behavior (8,10).

The beating motion of respiratory cilia follows an asymmetric pattern, comprising an active stroke and a recovery stroke (11). During the active stroke, the cilium stretches nearly straight to maximize its surface area against the PCL, propelling the ASL effectively. In the recovery stroke, the cilium bends significantly backward, resulting in less fluid propulsion. This coordinated motion, known as the cilia beat pattern (CBP), generates a net directional movement essential for MCC (12).

CFD simulations of MCC primarily focus on motile cilia, employing either direct cilia modeling or continuum models (13). Continuum models approximate the effect of cilia motion by introducing volume forces to simulate the CBP. While computationally efficient, this approach introduces non-physical effects, as seen in Roy (2013), where a viscoelastic “traction layer” was added at the ML-PCL boundary based on Lubkin’s (2007) method (14,15). Direct cilia modeling, on the other hand, captures the dynamics of cilium-fluid interactions more accurately but is computationally expensive. Blake’s (1972) pioneering model represented cilia positions using a truncated Fourier series based on experimental data (16). More recent approaches, such as Chateau et al. (2017), employed parametric curves to solve 1D transport equations for cilia positioning over time (17). Additionally, the immersed boundary method (IBM) was adapted by Quek (2018), representing cilia as linear and torsional elastic springs (18). Further advancements by Dillon (2007) and Yang (2008) introduced models that incorporated dynein motor dynamics and two-way fluid-cilium coupling (19,20).

Modeling the rheology of mucus adds another layer of complexity. Mucus exhibits non-Newtonian and viscoelastic properties, making mathematical modeling challenging. While earlier MCC models often neglected inertial terms in the Navier-Stokes equation (the Stokeslet method), more recent studies have emphasized their inclusion for improved accuracy (10,13). Notably, mucus’s shear-thinning behavior necessitates non-Newtonian rheological models, as assuming constant viscosity under shear stress leads to inaccuracies. A review by Vanaki (2020) summarized 17 CFD models of MCC, highlighting the predominance of Newtonian assumptions in earlier studies (13). Recent advancements, however, have increasingly adopted non-Newtonian fluid models. For instance, Chatelain (2017) and Barlett (2023) used the Carreau model, Sedaghat (2023) applied the 5-Mode Giesekus model from Vasquez (2016), and Shaheen (2023) utilized the Rabinowitsch model (4,6,21–23).

Contextually, this study introduces a two-phase fluid-structure interaction (FSI) simulation with direct cilim modeling, incorporating a non-Newtonian Carreau model for mucus rheology. A novel method for directly modeling the CBP is developed and evaluated within a two-dimensional framework. The prescribed displacement method, and the effects of non-Newtonian versus Newtonian mucus rheology are compared to establish a replicable two-phase FSI model of MCC.

## 2. Materials & Methods

### 2.1 Model setup

#### 2.1.1 Geometrical Model and Moving Mesh

Figure 1 shows the biological reference and the adopted geometrical model, consisting of two domains: the upper domain represents the area containing the ML (*Ω*_*ML*_), while the bottom domain represents the PCL (*Ω*_*PCL*_). According to the literature, the heights were set to 4 and 7 *μ*m for the ML and the PCL, respectively, as shown in Table 1 reporting the parameters’ values and the related reference. The length of the domain was arbitrarily set to 25 *μ*m to better visualize the effects in the fluids caused by the cilium motion. The cilium was located in the center of the domain in the horizontal direction, with its base attached to the bottom of the PCL (*Γ*_*L*_), representing lung tissue. The cilium height was set to 6 *μ*m and its diameter (thickness) was set to 0.2 *μ*m. A fillet was added to the top of the cilium with a 0.2 *μ*m diameter to represent lung cilia geometry more accurately. There is a discrepancy in previous models between if the cilia should penetrate the ML, or if it should beat entirely within the PCL, with the very top of the cilium being just lower than the PCL height when the cilium is vertically upright. The PCL height was adjusted to 7 *μ*m as to not allow the cilium to penetrate the ML and fit the assumption that the cilium beats entirely with the PCL.

**Figure 1:**
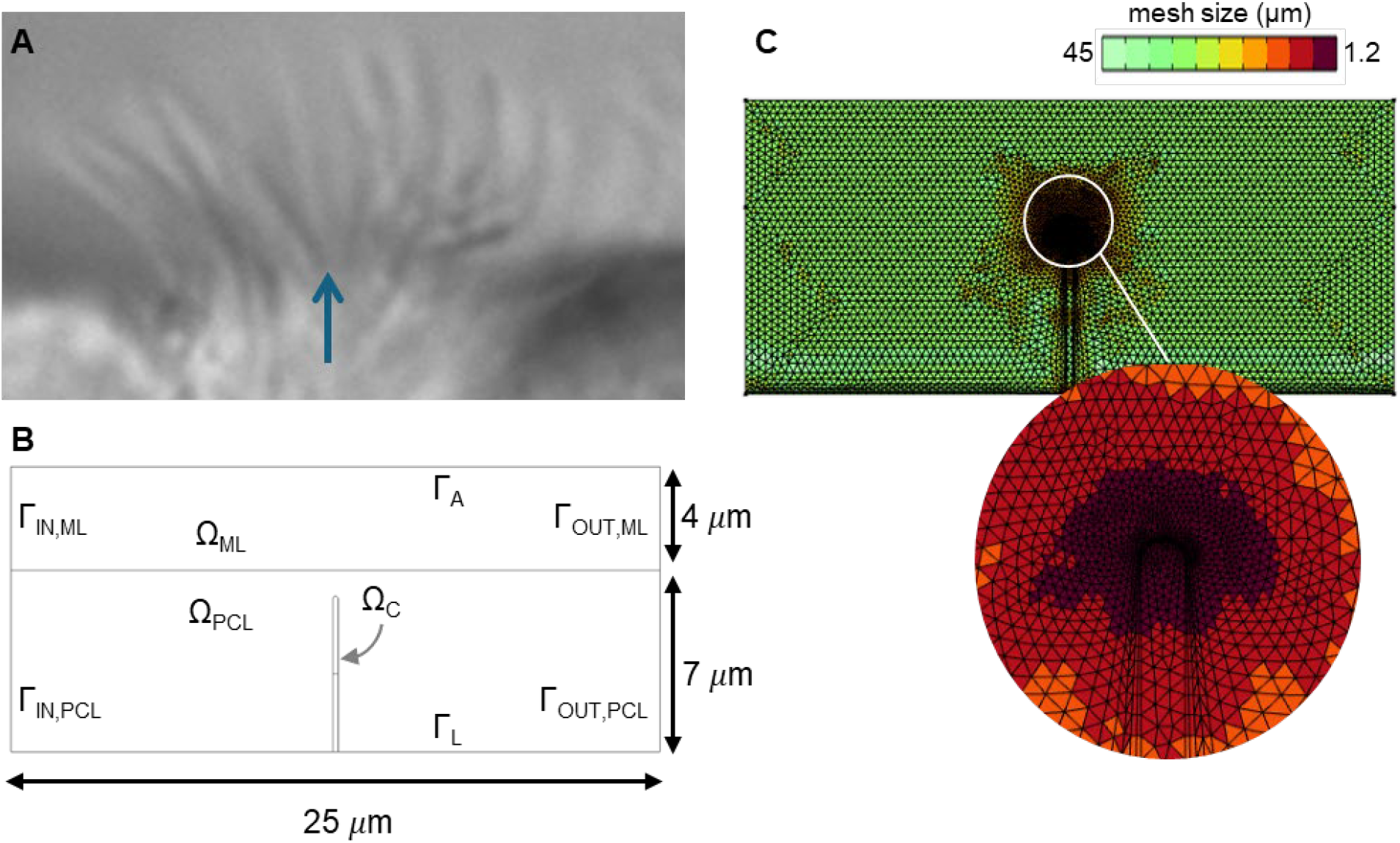
Cilium model development. **A**) One frame of the original (microscopy) data of the nasal swab. The blue arrow indicates the location of the cilium of interest and then manually analysed. **B**) Schematic of the adapted geometrical model, consisting of 2 fluid domains composing the airways surface liquid (Ω_ML_ and Ω_PCL_) and the cilium structural domain (Ω_C_). The inlet (Γ_IN,ML_ and Γ_IN,PCL_) and outlet (Γ_OUT,ML_ and Γ_OUT,PCL_) walls are prescribed fluid boundary conditions; while the top (Γ_A_) and bottom (Γ_L_) walls representing the interface with air and lung epithelium are prescribed slip and no-slip conditions, respectively. **C**) Final computational grid chosen following a mesh independence analysis, shown at rest phase (t=0). On the bottom, zoom on the mesh elements around cilium tip. The color legend represents the mesh element dimension.

**Table 1:** Model parameters.

| Parameter | U.M. | Value | Reference |
| --- | --- | --- | --- |
| Cilium Height | $\mu\text{m}$ | 6 | (36) |
| Cilium Diameter | $\mu\text{m}$ | 0.2 | (37) |
| Cilium Elasticity Modulus | MPa | 0.764 | (38) |
| Cilium Beat Frequency | Hz | 3 | (39) |
| PCL Height | $\mu\text{m}$ | 6 | (40) |
| PCL and ML Density | $\text{Pa} \cdot \text{s}$ | 1000 | (15) |
| PCL Viscosity | $\text{Pa} \cdot \text{s}$ | 0.001 | (15) |
| ML Height | $\mu\text{m}$ | 4 | (41) |
| ML Viscosity (Newtonian) | $\text{Pa} \cdot \text{s}$ | 0.1 | (42) |
| ML Zero Shear Rate Viscosity | $\text{Pa} \cdot \text{s}$ | 302 | (4,43) |
| ML Infinite Shear Rate Viscosity | $\text{Pa} \cdot \text{s}$ | 0 | (4,43) |
| Relaxation Time | s | 8440 | (4,43) |
| Power Index | N/A | 0.375 | (4,43) |

A triangular moving mesh was used, with a final number of 10’952 elements obtained from grid independence analysis (see Supplementary Materials), with a final minimum element length of ∼1.2 *μ*m. Figure 1-C shows the larger concentration of elements around the cilium tip. With the given maximum element length, a maximum fluid velocity of 100 *μ*m/s and a maximum time step set to 0.001 s were set, verifying the Courant–Friedrichs–Lewy condition through:

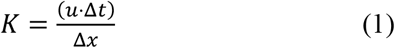

Contextually, this provided a value of 0.2137 - lower than the usual recommended maximum value of 1 (24).

#### 2.1.2 Boundary Conditions

The bottom (*Γ*_*L*_) and top (*Γ*_*A*_) walls represent the PCL interface with lung tissue and the ML interface with air, respectively. They were both modeled as fixed boundaries, constraining the base of the cilium in the bottom wall *Γ*_*L*_. To represent the low shear stress between the ML and the air, a slip condition was imposed at the top wall *Γ*_*A*_. Conversely, a no-slip condition was imposed at the bottom wall *Γ*_*L*_ to represent the shear between the lung tissue and PCL. Concerning the lateral walls, the left side of both domains was set as inlet (*Γ*_*IN*_) while the right side of both domains *Ω*_*ML*_ and *Ω*_*PCL*_ was set as outlet (*Γ*_*OUT*_), implying a net flow from left to right. The initial system was set at rest, namely the initial velocity of the ML and PCL, was set to 0 *μ*m/s.

### 2.2 Model Equations

#### 2.2.1 Airway Surface Liquid fluids

The PCL is modeled as Newtonian fluid, assuming incompressible and laminar flow. The CFD simulation describes this flow mathematically using the Navier-Stokes equation and the continuity equation in the following forms:

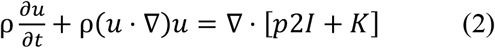

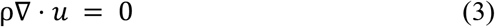

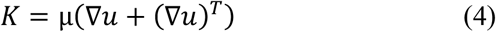

In the above equations, ρ is density, *u* is velocity, *t* is time, *p* is pressure, *I* is the identity tensor, *K* is the stress tensor, and μ is the dynamic viscosity. These parameters and the above equations will determine how the PCL interacts with the system.

The ML is modelled instead as a non-Newtonian fluid assuming incompressible and laminar flow and is modeled with the same equations as the PCL. However, there is the significant dependence of the ML’s viscosity on its shear strain rate. Therefore, Equations (2-4) for the PCL will still be used, but additional equations representing the viscosity will be added based on the non-Newtonian Carreau model:

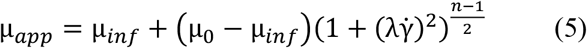

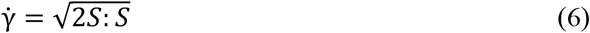

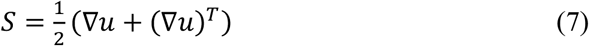

In the above, μ_*app*_ is the viscosity considering shear thinning, μ_*inf*_ is the infinite shear rate viscosity, μ_0_ is the zero shear rate viscosity, λ is the relaxation time, *n* is the power index, 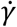 is the shear rate. and *S* is the strain tensor.

#### 2.2.2 Cilia-ASL interaction

FSI models the effect that the cilia’s motion has on the fluid flowing around it, coupling the velocity of the two. This interaction is governed by the following equation, representing the force exerted by the fluid onto the structure (cilium):

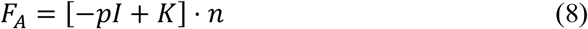

where *p* is pressure, *I* is the identity matrix, *K* is expressed in Equation 4, and *n* is the normal vector.

### 2.3 Cilia Displacement Prescription

As previously mentioned, MCC is driven by cilium motion. In this model, MCC is modeled as a cyclic prescribed displacement given to points along the cilium, giving the cilium a CBP that drives the fluid motion.

#### 2.3.1 Displacement Spline

The position over time of the cilium was extracted from open-source data (25). The data was processed using ImageJ, in which high speed video frames of a CBP were manually processed frame by frame by recording the image coordinates of the cilium’s origin, the cilium’s tip, and points along the cilium throughout its beat cycle. The data is then fed to Python where it is scaled to have an initial position of 0 so that its movements are relative to its starting position and scaled to represent the true length of the cilium (in this case to the standard length of lung cilia).

A B-spline was made to represent the position over time of the cilium. The B-spline uses the knots, t, the spline coefficients, c, and the B-spline degree, k, to produce a B-Spline object. The mathematical representation of a B-Spline in summation from is given here:

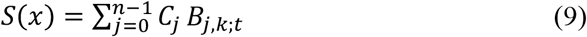

where B are spline basis functions. The Cox–de Boor recursion formula is used to construct the B-spline:

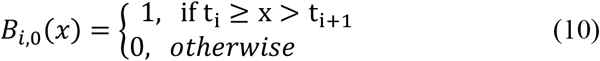

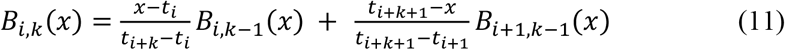

The knots, spline coefficients, and B-spline degree (t, c, and k) were found by introducing the dataset retrieved experimentally and giving t, c, and k for a chosen smoothing parameter s, employing the python package scipy.interpolate. The smoothness is given by a variation of the squared error which can be skewed to weigh certain parts of the data more based on the selection of the w vector:

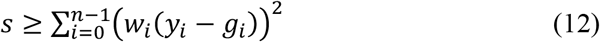

where g is the smoothed interpolation of y.

Now, with the experimental data retrieved and a chosen s-value, a B-spline can be created with a desired smoothness. The value for s is determined by the calculation of the r2 value and the mean squared error (MSE). With these two metrics it can be ensured that the spline representation is accurate to the desired degree, while manual inspection allows for determining if the curve is smooth enough so that overfitting is not an issue. Overfitting may also lead to convergence issues in simulations due to the sharpness of the velocity path if the spline is not smoothed. Selected splines for different smoothness values, including the chosen value of *s* = 15, are shown in Figure 2-B. The spline used in simulations gave an r^2^ score of 0.970 and an MSE of 0.109.

**Figure 2:**
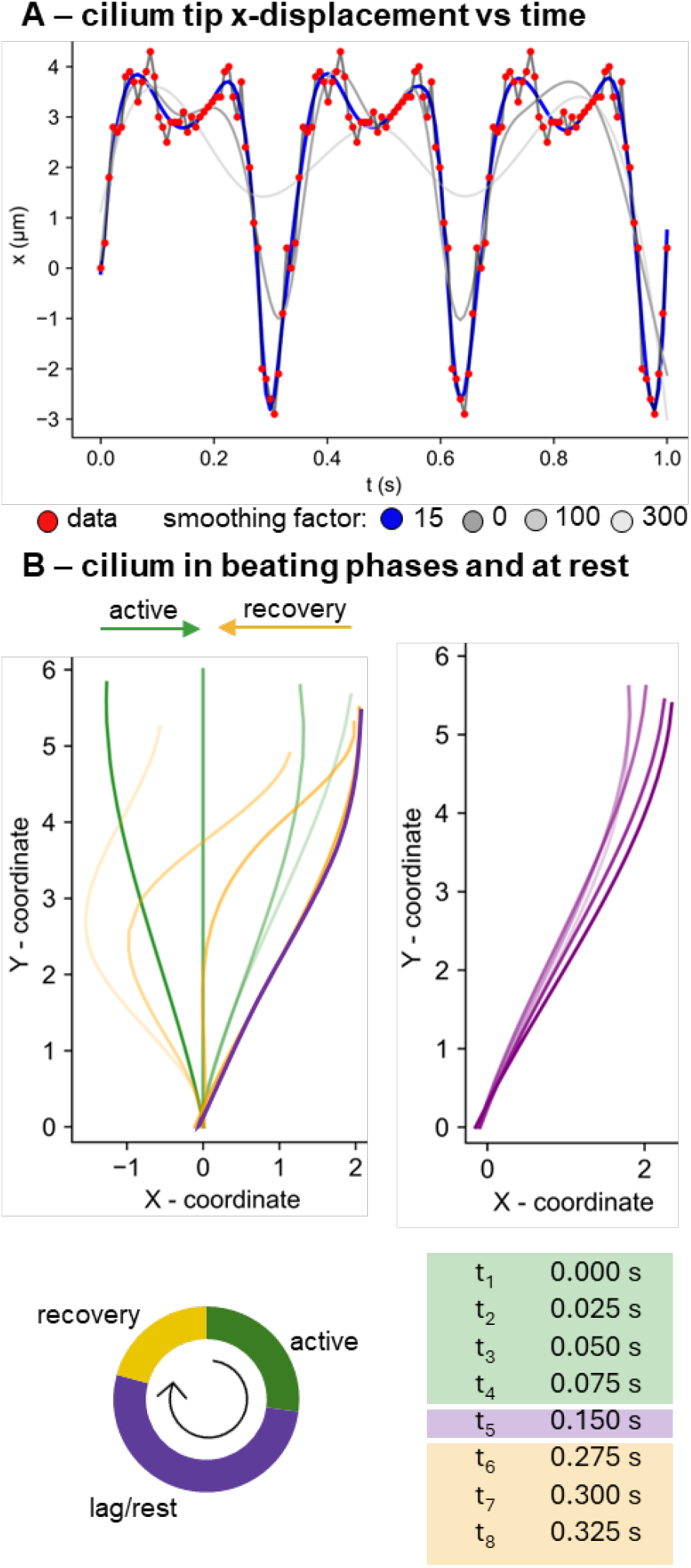
Cilium displacement. **A**) Cilium tip displacement in the x-direction along 3 beating cycles. The data were interpolated with B-splines smoothed with incremental factors, as shown for example in this panel. The blue line indicates the final chosen smoothing factor (=15). Red dots are the measured data used for the B-spline fitting. **B**) displacement in the x and y directions of the cilium during the beating cycle (left panel). Starting from the rest configuration (indicated with 0), the cilium beats first forward (represented with yellow shades) and then backward (with green shades) in the active and recovery phases, respectively. The considered rest position (lag) is the averaged of the rest configurations (in violet shades) showed in the right panel.

#### 2.3.2 Mid-point Spline

Cilia deform in nonlinear ways, making it difficult to find the exact midpoint or desired point along the cilium at different frames to label it to use for a prescribed displacement. This makes adding a prescribed displacement to a section of the cilium below the tip more difficult.

To find the desired points along the cilium, another B-spline is created to represent the two-dimensional profile of the cilium midline spatially, rather than a point on the cilium over time as with the previous spline. Once the B-Spline representing the cilium shape is retrieved, very small steps in the horizontal direction are taken, and the position along the spline is calculated. The total change in cilium length at each iteration is then calculated using the general displacement equation given below and then added to the total.

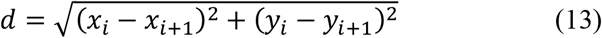

Once the total length of the cilium at a specific frame is known, the code will iterate through the spline again and print the coordinates of the midpoint (or any desired point). This is then repeated for every frame, and with this information, the same outlined process can be used to find a spline representing the displacement over time at any point on the cilium. The final B-spline representing the displacements of points on the cilium was then imposed as prescribed displacement in the FSI solver, as presented in the following section.

#### 2.3.3 Solver settings

The simulations were solved in COMSOL Multiphysics (v. 6.1, Stockholm, Sweden). A reinitialization parameter was set to 7 *μ*m/s to help with convergence, approximating the velocity caused by the cilium. A relative tolerance of 0.001 was given to the stationary solver. A relative tolerance of 0.3 was imposed to the time-dependent solver and an absolute tolerance in the form of a scaled tolerance was also prescribed to the time-dependent solver and was set to 0.05. A segregated solver was used, selecting Constant (Newton) method for all segregated solvers except for the segregated FSI solver which was set to Automatic highly nonlinear (Newton) to help with convergence. A moving mesh was used throughout all domains in the present simulations to ensure the mesh remains finest in the areas of highest fluid velocity as the prescribed displacement of the cilium causes these areas to move.

### 2.4 Analysed scenarios and computational MCC indexes

To compare the impact of computational settings on the mucus flow, we considered 4 different scenarios. Specifically, while keeping the PCL Newtonian rheology, we assessed the impact of prescribing non-Newtonian properties to the ML. Similarly, we compared the effect of performing a one- or two-point displacement prescription to the cilium. The analysed scenarios are defined as follows: case 1 (“C1”), where Newtonian ML and one-point displacement were set; case 2 (“C2”), where non-Newtonian ML and one-point displacement were imposed; case 3 (“C3”), where Newtonian ML and two-point displacement were given; and case 4 (“C4”), where non-Newtonian ML and two-point displacement were prescribed.

Besides analysing the changes in Von Mises Stress, velocity field, and volume fraction between the models, we also propose to quantify the overall effect of the computational prescriptions on the MCC. It is known that, in vivo, MCC is typically measured through tracking the motion of inhaled tracers passively transported by the ciliary action (26) Here, for the sake of comparing the computational models, we proposed two computational MCC indexes. First, we computed the percentage of normalized flux (*nF*) contributions in the active, lag and rest phases. For each phase, it was obtained as the ratio between the signed flux of the x-component velocities of the related phase and the sum of the absolute values of the phase fluxes. This index allowed to observe the contribution and directionality of the mucus flow during the phases of the cilium beating cycle. The individual fluxes were computed for each simulation time-step as the product between the median x-component velocities and the arc length (4µm) at the center of the ML domain (Ω_ML_). The second index, based on the normalized flux *nF*, is the net mucus flux (*nMF*) which is the sum of the three *nF*. The latter indicate the flux amount that is measured at the end of the cilium beating cycle.

The velocity distributions were tested for normality through the Kolmogorov Smirnov test, with a significance level of 5%. Following the result, non-parametric tests were adopted, and the values were reported as median and the inter-quartile range (IQR). The comparison among groups of the velocity distributions were tested through the Kruskal-Wallis ANOVA test. The analyses were performed in MATLAB (R2022b).

## 3. Results

### Cilium structural deformation

Figure 3 shows the Von Mises Stress induced by the PCL domain on the cilium during the beating cycle, from active phase onset (t_1_), through the lag phase (t_5_), and then to the end of recovery phase (t_8_). It can be observed that the major variations of the structural stress distributions arise from the displacement prescription, being the structural stress patterns pairwise similar in the cases C1-C2 and C3-C4. The maximum Von Mises stress value was found in C4 during the transition from the recovery to active phase, i.e. at t_8_, with a value of 224.7 kN/m^2^. The respective value in the cases C1-3 was, respectively, 5.5, 4.7, and 214.3 kN/m^2^. The minimum Von Mises Stress at the same time instat was found to be, in order: 0.08, 0.098, 1.323 and 0.78 kN/m^2^ for C1-C4.

**Figure 3:**
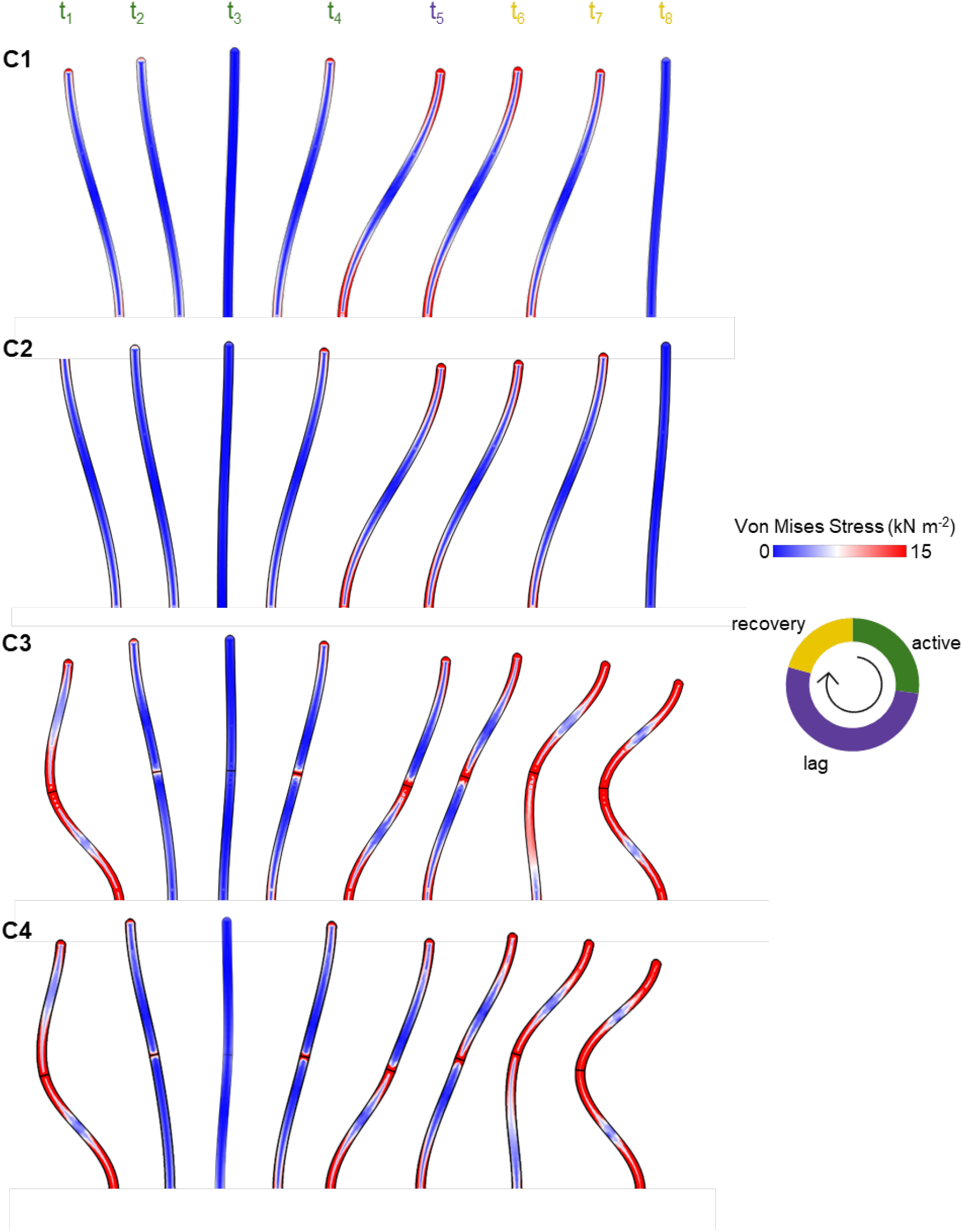
Cilium structural deformation. Von Mises Stress induced by the periciliary liquid on the cilium at the representative time instants: in the active (t_1_ – t_4_, in green), lag (t_5,_ in purple) and recovery phases (t_6_ – t_8_, in yellow), with non-newtonian ASL rheology and two-point prescribed displament.

### Influence of ASL rheology

To assess the influence of ASL rheology, we first observed the velocity magnitude at two selected time instants, namely the end of the active phase (t_4_) and the end of the recovery phase (t_8_) (Figure 4A). By qualitatively comparing the cases C1 (Newtonian ML) and C2 (non-Newtonian ML), it resulted that ML rheology influences the fluid dynamics in PCL. More in detail, the non-Newtonian model (C2) led to an approximately two-fold increase in the maximum velocity of the PCL for both the active and recovery strokes, as compared to the Newtonian model (C1). Moving to a more quantitative comparison of the ML fluid dynamics (Figure 5), limited and statistically non-significant differences were found between the velocities in the Newtonian (C1) and the non-Newtonian (C2) cases, both presenting the one-point cilium displacement prescription. Specifically, from Figure 5-A and -B, the distribution of the x-velocities in the ML above the cilium were alike in the lag and recovery phases, as supported by the resulting statical non significance. Slight differences, non-significant though, were found in the active phase, where C2 presented both higher maximum and lower minimum x-velocities.

**Figure 4:**
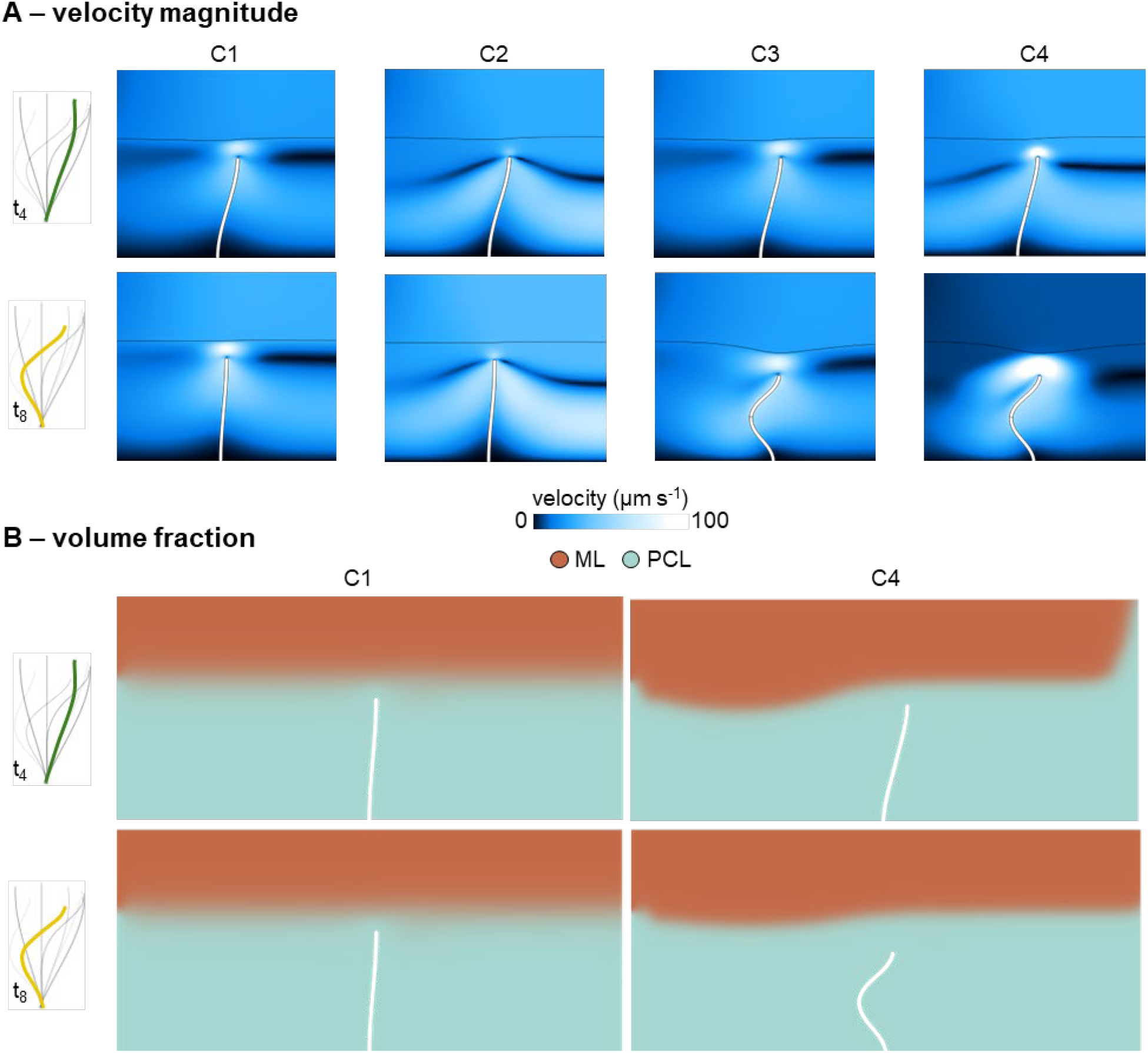
Airway surface liquid (ASL) velocity and volume fraction results. **A**) Countour maps of ASL velocity magnitude evaluated at t_4_ (end of active phase) and at t_8_ (end of recovery phase) in the four scenarios C1-4. While minor changes are visible in the velocity of mucus layer (ML) at t_4_, substantial differences are found between the recovery ML speed in C4. **B)** Contour maps of the volume fraction change in cases C1 and C4 at the two considered time instants. C4 non-Newtonian model and two-point displacement prescription induces larger modifications in the ML and periciliary layer (PCL) distributions.

**Figure 5:**
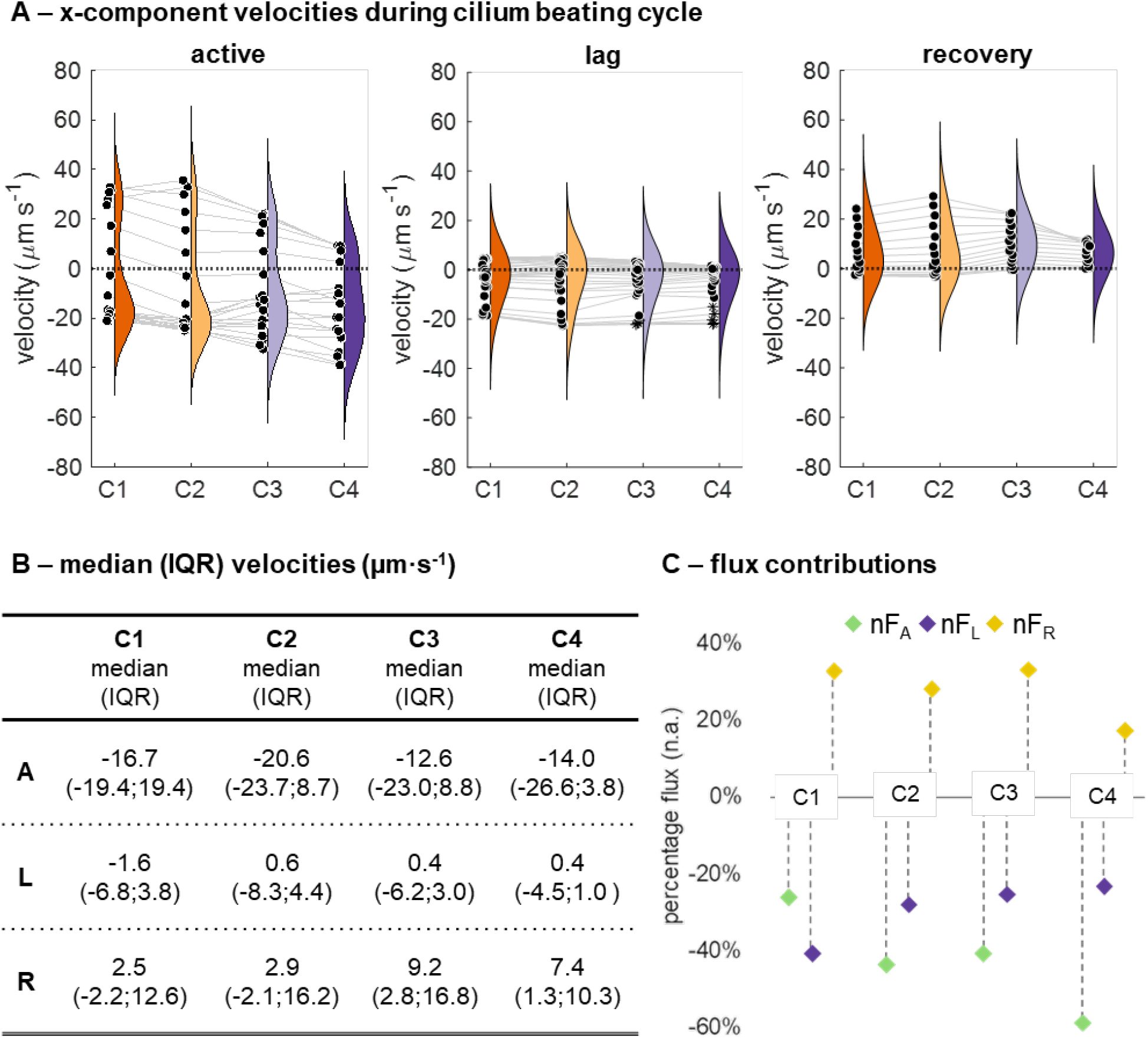
Mucus layer (ML) velocities during the beating phases. **A**) On the top, for each time step of the beating cycle, the median of the x-component velocities, extracted from the mesh-interpolation in the ML above the cilium, are computed and represented in the violin plots, subdivided for active, lag, and recovery phases. The grey lines connect the matched time-steps in the four analysed cases. While negligible variations are found during the lag phase, the x-velocities have lower median in C4. **B**) Median and inter-quartile range (IQR) velocity values for the three phases and the four scenarios. **C**) Percentage flux contributions of the active, lag, and recovery phases, normalized to the total sum of the absolute value of the individual flux contributions, computed on the time-step x-velocity values. It is observed that the maximum flux contribution in the active phase *nF*_*A*_ is found in C4. A: active; L: lag; R: recovery; nF: normalized flux.

### Influence of cilium movement

Analogous observations emerged in comparing C1 and C3, where the impact of one-point or two-point displacement prescription was imposed in a Newtonian ASL model. From Figure 4-A, it emerges that negligible differences in velocity magnitude in the ASL were determined by the cilium movement prescription methodology at the time instant t_4_. However, while the velocity field was not quantitatively affected (no statistically significant differences were found), it is worth noting that the PCL-ML interface was affected by the cilium deformations occurring in the recovery time instant t_8_.

However, for the non-Newtonian two-point, there is only an increase for the active stroke, as there is a large decrease in velocity on the recovery stroke. In these graphs, the coupling between the upper PCL and ML for the non-Newtonian models is also apparent (where the horizontal line at 7 *μ*m corresponds to the PCL-ML interface).

### Selected model: non-Newtonian ML and two-point displacement

The case C4, with non-Newtonian ML rheology and two-point displacement prescription, presented the major changes compared to the other models. Merged observations from the previous sections can be made. In the active phase (t_4_) (Figure 4-A), case C4 qualitatively presented similar velocity field patterns in the ML, as supported by the statistical non-significant difference found (chi-square: 4.04, p-value: 0.26). The major qualitative changes can be observed in the PCL, with limited relevance given the absence of other cilia in the model. This aspect is further discussed in the following section. The recovery stroke found in C4 resembled real cilia motion more accurately with a two-point prescribed displacement, as expected, as it allows for the capturing of the bending of the cilium during the recovery stroke. Unexpectedly, the two-point prescribed displacement led to non-significant differences in the velocity distributions during the beating cycle (Figure 5A,B). Quantitatively, by considering all the time steps of the cilium beating cycle, C4 presented a median velocity of -14.0 µm·s^-1^, 0.4 µm·s^-1^, and 7.4 µm·s^-1^ in the active, lag and recovery phases, respectively. No statistically significant differences were found compared to the other cases neither in the lag (chi-square: 2.07, p-value: 0.56), nor in the recovery (chi-square: 3.01, p-value: 0.39) phases.

As for C3, the bending of the PCL-ML interface is visible in Figure 4-A. The countour map of the volume fractions (Figure 4-B) indicates that the ML fraction is not confined in the upward region (above the interface construction line), in both the active and the recovery phases of C4. It is worth noting that the volume fraction in the upper part of the model (above the interface line) does not differ in the active phase in the other models (C1-3, see also Supplementary Figure S2). The non-Newtonian ML rheology imposed in C2 leads to volume fraction changes in the region below the interface line, but only in the recovery phase. It can be concluded that the overall contribution of the non-Newtonian rheology and the two-point prescription (model C4) provides a more realistic representation of the multiphase interaction.

Last, Figure 5-C shows that C4 has a larger negative *nF* in the active phase (−59%), similar *nF* in the lag phase (−23%), and minimum *nF* in the recovery phase (18%). Regarding the net flux in the ML, *nMF* increases from the lowest value for C1 (33%), to the largest value (64%) for C4. *nMF* in C2 and C3 was respectively 42% and 33%, indicating that the non-Newtonian rheology contributed largely to the changes of the mucus flux. Therefore, the combination of non-Newtonian ML and two-point displacement gave a non-trivial modification to the *nMF*, confirming that the combination of these two modelling aspects leads to non-linear modification in the ML fluid field.

## 4. Discussion

In this study, we developed a two-phase FSI model of MCC, imposing displacement on the cilium. The primary objective was to create a replicable and modular model, capable of easily adapting fluid properties and cilium motions. Given the impact of computational settings and decisions on the flow results in the mucus layer (ML), it was vital to assess the influence of these choices. This prompted a comparison of ML rheology (Newtonian vs. non-Newtonian) and cilium displacement prescriptions (specifically, how cilium motion determines the cilia beat pattern, CBP).

Recent advancements in direct cilia modeling for MCC have emphasized incorporating the shear-thinning effects of non-Newtonian ML (13). Our findings demonstrate improved coupling between the ML and the upper PCL when a non-Newtonian model (e.g., Carreau, C2, and C4) is applied (see Figures 4 and 5). In contrast, Newtonian models (C1 and C3) maintained a distinct ML-PCL interface boundary. This coupling in the non-Newtonian model resulted in increased ML velocity and enhanced MCC rates.

Comparing our results with literature (15,21,27–32), we observed that average ML velocity typically ranged between 10 – 92 *μ*m/s (with the majority in the 35-45 *μ*m/s range) for Newtonian, viscoelastic, or non-Newtonian simulations, and 39.2 *μ*m/s from in vitro experiments (28). In our simulations, the ML velocity magnitude ranged from approximately 17 to 39 *μ*m/s during the active stroke, and between 4 and 26 *μ*m/s on the recovery stroke, aligning with these reported ranges. Modaresi and colleagues further validated this conclusion, showing that non-Newtonian scenarios (power law and thixotropic fluids), generally leads to higher MCC rates compared to Newtonian scenarios, consistent with our computational MCC indexes. Furthermore, Chatelain (2016) simulated healthy cilia using non-Newtonian ML rheology, finding velocities of approximately 34 – 38 *μ*m/s, in strong agreement with the two-point non-Newtonian model in our work (21).

We also explored the impact of cilium motion and CBP on the ML, employing a novel method of prescribing displacement at specific points along the cilium. Although this approach mirrors techniques in adjacent fields (e.g., for algae cilia) (33), challenges arose due to the aggregation of experimental cilia data. Tracing displacement for more than three points on the cilium (including the epithelium attachment) often resulted in unrealistic motion. Our comparison of one-point and two-point prescribed displacements revealed that a two-point approach (tip and mid-cilium) produced more realistic interactions between the PCL and ML, as evidenced in Figure 4-B. The advantage of the here-presented modelling approach is that the cilium motion pattern, driving the fluid dynamics, can be directly derived from other modelling systems, such as systems biology, molecular dynamics, and agent-based model approaches (32,34,35) and then directly imposed to the continuum FSI model, providing an innovative versatile platform for FSI modelling of MCC.

While the current FSI methodology represents a significant advance, several improvements are needed for greater accuracy. First, enhanced experimental data could enable finer displacement prescriptions, improving the fidelity of cilium movement. Extending the model to three dimensions would allow for simulating metachronal waves—coordinated cilia motions—that are critical for MCC (13). Additionally, incorporating surface tension effects, as Quek (2018) suggested, could further enhance ML velocity predictions in three-dimensional frameworks (18). Nevertheless, earlier studies (27,30), found minimal impact of surface tension on two-dimensional ML models at levels up to 10 N·m^-1^, highlighting the need for further exploration. Last, further changes need to be made for this model to accurately represent MCC. Specifically, changes to simulation parameters should be modified to reduce numerical inaccuracies, and additions to the setup of the simulation should be made to create an environment that more closely represents the cilia environment lining lung epithelium.

This study successfully introduced a two-phase FSI model of MCC, evaluating non-Newtonian ML rheology and an innovative cilium displacement method. The findings underscore the importance of non-Newtonian rheology and multi-point prescribed displacements (at least two) in achieving accurate MCC representations. Our computational MCC indexes, *nF* and *aML*, provide valuable metrics for understanding the influence of computational choices on model outcomes. The adaptability of the novel displacement techniques opens pathways for multi-scale, multi-methodological approaches, facilitating investigations into ciliary disorders from cellular mechanisms to continuum dynamics.

## Acknowledgments

Monika Colombo and Subburraj Karupppasamy acknowledge support from Aarhus University.

