## Supplementary Materials for "A Two-Phase Fluid Structure Interaction Model of Mucociliary Clearance Driven by Cilium"

# of

### S.1 Grid independence analysis

**Table S1: Parameters of the grid independence analysis.** By employing the physics-controlled mesh generation in COMSOL, the maximum pressure and maximum velocities are computed at the outlet section at the beating time-step of 0.65 s. Convergence was not reached for the coarse and extra-fine cases (indicated in red).

| COMSOL Setting | Minimum Element Length (µm) | Maximum Element Length (µm) | Average Pressure (Pa) | Maximum Pressure (Pa) | Average Velocity (µm/s) | Maximum Velocity (µm/s) |
|---|---|---|---|---|---|---|
| Coarse | 0.0135 | 0.708 | N/A | N/A | N/A | N/A |
| Normal | 0.0122 | 0.598 | | 0.042 | | 16.8 |
| Fine | 0.0106 | 0.456 | | 0.050 | | 17.0 |
| Finer | 0.0112 | 0.225 | | 0.065 | | 18.9 |
| Extra Fine | 0.0109 | 0.106 | N/A | N/A | N/A | N/A |

### S.2 Statistical analysis: velocity distribution comparison

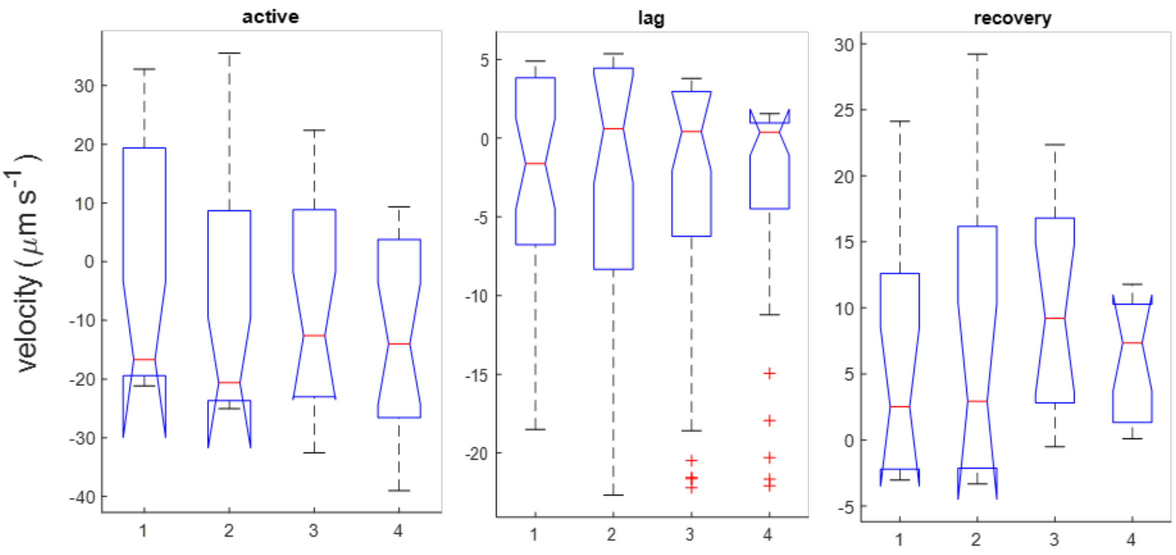


**Figure S1: Results of the group normal distribution comparison.** Results of the Kruskal-Wallis test performed for the velocity distributions in the case of active, lag and recovery phases for the four investigated scenarios. No statistically significant differences were found among the groups.

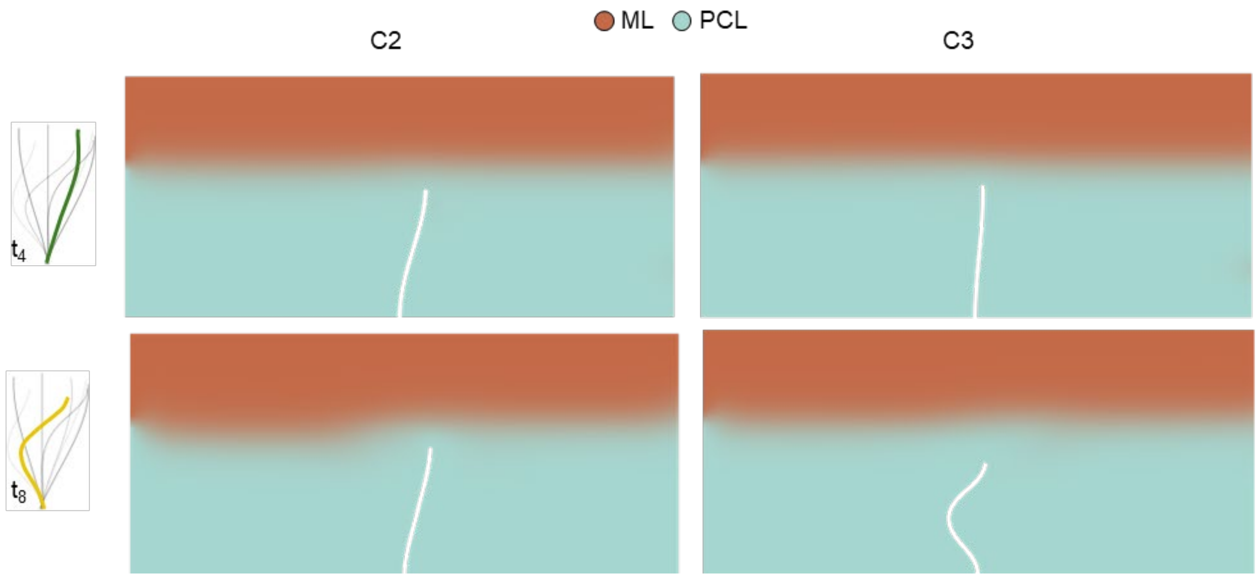


**Figure S2: Volume fraction changes.** Contour maps of the volume fraction change in cases C2 and C3 at the two considered time instants. ML: mucus layer; PCL: periciliary layer.
